# Multiphase capillarics for structurally pre-programmed sequential and parallel operations of high and low cohesion liquids

**DOI:** 10.64898/2026.09.08.749925

**Authors:** Geunyong Kim, Lan Anh Huynh, Molly L. Shen, Yonatan Morocz, Andy Ng, David Juncker

## Abstract

Capillaric circuits (CCs) enable pre-programmed liquid handling through self-filling and passive valving governed by capillary forces, eliminating the need for external pumps and actuators. However, CCs optimized for high cohesion liquids (HCL) such as water are intrinsically unsuitable for programmed flow control of low cohesion liquids (LCL) such as oils and solvents because the high surface free energy needed for capillary flow of HCL results in uncontrolled, complete wetting by LCL. Here, we introduce a library of multiphase components including phase-to-phase valves (P2PV), hydro-pneumatic relays (HPR) and multiphase domino valves (MDVs) that collectively enable CCs to concomitantly process HCLs and LCLs. P2PVs use a pre-filled HCL to valve immiscible LCLs (e.g. oil) by confining the LCL, and upon triggering, hydraulically entrain it. To prevent uncontrolled mixing between miscible LCLs (e.g. ethanol) and the HCL, HPRs with an air gap and air waste are added to the P2PV. MDVs are further added and enable preprogrammed, sequential delivery of LCL and HCL by multiphase microfluidic chain reactions. Multiphase liquid processing is applied to automated, on-chip lipid nanoparticle (LNP) manufacturing, illustrating the potential of multiphase CCs.

## Introduction

Multiphase liquid handling is used in a wide range of biomedical applications and includes an aqueous phase with either immiscible phases (e.g., oil and aqueous two-phase systems; ATPS) or miscible phases (e.g., alcohols and surfactants) that are introduced simultaneously or sequentially. Processes include solvent exchange (e.g., nucleic acid extraction^1,2^), mixing (e.g., LNP fabrication^3–6^), droplet generation and emulsification (e.g., formation of reaction compartments or microparticles for high-throughput analysis and digital assays^7–10^), phase-based separation (e.g., patterning and partitioning of cells and biomolecules using ATPS^11^) and sealing (e.g., oil capping in digital assays^12,13^). Water is a high cohesion liquid (HCL) that requires high free energy surface to promote surface wetting and the spontaneous filling of microchannels, while oils and solvents are low cohesion liquids (LCLs) that readily wet most surfaces, even those with comparatively low surface free energy^14^. Compared to single-phase aqueous systems, multiphase systems exhibit interfacial phenomena that can give rise to capillary pressure at the interface between immiscible phases, and in miscible systems, Marangoni flows driven by surface tension gradients, along with dynamic changes in wettability and viscosity during mixing. Precise control over these coupled hydrodynamic and interfacial effects is essential for reliable operations of multiphase systems.

Many microfluidic approaches have been developed for handling and automating multiphase liquid operations^15,16^, using pumps^4,7,8^, and centrifugal actuation^9^ with advanced, computer-programmable peripheral control systems. However, the reliance on peripheral control constitutes a barrier to broader adoption.

Capillary microfluidics enable self-powered liquid delivery and autonomous flow control and exist in two main formats including: (i) Paper-based capillary microfluidics (also known as microfluidic paper-based analytical devices; µPADs) that are widely used as lateral flow assays (LFAs) and in applications such as wearable devices^17–19^, and (ii) Microchannel-based capillary microfluidics^20–25^ also termed capillarics in analogy to electronics^20–22^. Originally fabricated via cleanroom microfabrication as well as soft lithography, capillaric circuits (CCs) can now be 3D-printed, offering a more accessible and rapid route to device prototyping and manufacturing^26,27^. CCs structurally encode algorithms and autonomously execute sequential liquid delivery steps, which has been demonstrated for 300 steps using a microfluidic chain reaction (MCR)^23^.

Autonomous liquid operation in CCs requires spontaneous capillary flow with geometric control over flow stop and start using capillary valves^20,21^. Stop valves (SVs) are formed by abrupt expansions from a shallow, narrow microchannel to a larger, deeper one and pin the advancing liquid front at the edge of the junction between the two channels. Flow can also be controlled using retention burst valves (RBVs) and capillary retention valves (CRVs) that act on the back, receding end of the liquid plug, and resist liquid drainage from the downstream microchannel by generating negative capillary pressure. RBVs and CRVs are distinguished by whether their burst pressure is exceeded by the driving capillary pressure at the filling front: valves that burst under the filling front pressure are classified as RBVs, whereas those that do not are CRVs. In CCs, liquid reservoirs are connected to the main channel through shallow and narrow channels termed functional connections, which form SVs at their junctions with the larger main channel. Upon reservoir filling, these SVs stop the advancing liquid front and confine the liquid within the reservoir. Subsequent filling of the main channel establishes a continuous hydraulic connection between the reservoir and the main channel and, by extension, the capillary pump at its downstream end, driving structurally programmed sequential delivery of liquids from the reservoirs^21,23^.

Autonomous operation and flow control in CCs rely on adjusting the surface free energy of the microchannels so that a liquid droplet exhibits a contact angle of ∼30–60° (an empirically determined range) when measured on a flat surface with the same surface chemistry^20^. For an HCL such as water (surface tension *σ*_water_ = 72.8 mN/m), a comparatively high surface free energy is required to create hydrophilic surfaces. On the same surface, however, LCLs such as oils (e.g., *σ*_hexadecane_ = 27 mN/m) and alcohols (e.g., *σ*_ethanol_ = 22 mN/m) will have contact angle below the 30° threshold, and sometimes, as low as 0°. While LCLs can also spontaneously fill CCs with high surface free energy, the stop valves become dysfunctional, causing uncontrollable spreading. Conversely, if the surface free energy is lowered and the surface made hydrophobic, then the contact angle of an HCL will be > 60° (and often > 90°), and the HCL will no longer spontaneously fill the microchannels. Thus, the operational contact-angle window cannot be satisfied simultaneously for both LCLs and HCLs in conventional CCs. Spatial patterning of regions with high and low surface free energies could, in principle, address this limitation, but it introduces substantial complexity in fabrication^28^ and limits flexibility in CC designs. Consequently, CCs have thus far been limited to single liquid phase (and overwhelmingly aqueous phases) handling.

Here, we introduce a multiphase capillary valving architecture and a library of components that use an HCL as a control element to confine, valve, and actuate LCLs, and to enable the design and fabrication of 3D-printed multiphase CCs that can concomitantly process HCLs and LCLs. We further consider LCLs that are miscible and immiscible with a reference HCL. Using water as the reference HCL, immiscible LCLs include oils whereas miscible LCLs include solvents such as methanol, ethanol, isopropanol, etc., as well as water with surfactants. We further introduce air gaps and design architectures that allow the actuation of miscible LCLs while preventing unwanted mixing between LCLs and HCLs. We use this library of multiphase capillaric components to design an autonomous, self-contained LNP formulator. Upon loading the LCLs and HCLs, and connecting a paper pump, a multiphase MCR CC sequentially delivers, dilutes, mixes, displaces the various phases to fabricate functional LNPs.

## Results and discussion

We propose multiphase capillary valving architecture and a library of capillaric components including phase-to-phase valves (P2PVs), hydro-pneumatic relays (HPRs) and multiphase domino valves (MDVs), for coordinated multiphase liquid handling, Fig. 1a-c. The CCs use microliter-scale liquid reservoirs, with capacities typically ranging from a few to ∼10 µL depending on the application. For the valving and flow control of LCL in such reservoirs, P2PVs use HCLs to physically confine and hydraulically actuate them, thereby enabling structurally preprogrammed retention and delivery of LCLs. An HPR is added to a P2PV to isolate the LCL from the actuating HCL, thus preventing premature mixing of a miscible LCL and the HCL. HPRs include both an air gap and an air waste. The air gap initially disconnects the miscible LCL reservoir from the HCL in the main channel, and upon actuation, the air is directed into the waste, connecting the LCL reservoir to the main channel for delivery by hydraulic entrainment of the LCL. MDVs, supplemented with an HCL that isolate the LCLs, replace conventional capillary domino valves (CDVs)^23^, allowing integration of LCL reservoirs into the MCR. MDVs enable the sequential delivery of arbitrary combination of HCLs and LCLs, thus upgrading the MCR into a multiphase system compatible with LCLs. A multiphase MCR CC incorporating a staggered herringbone micromixer is used for automated on-chip fabrication of LNPs, Fig. 1d.

**Figure 1.**
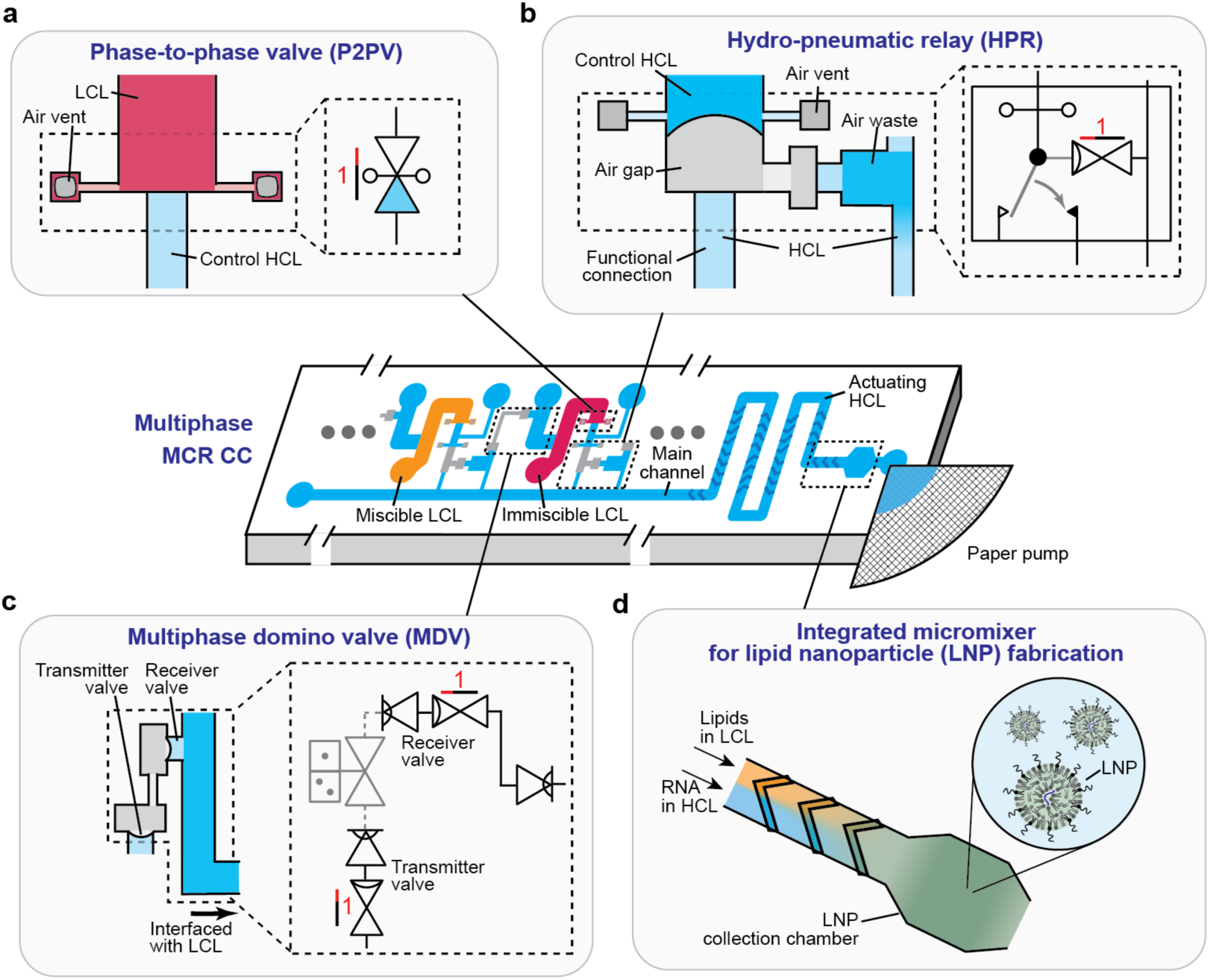
Library of multiphase control components for concomitant handling of high- and low-cohesion liquids. (a) A phase-to-phase valve (P2PV) uses a control high-cohesion liquid (HCL) to hydraulically control an LCL. (b) Hydro-pneumatic relay (HPR) with an air gap and an air waste. HPRs are applied downstream of P2PVs and prevent premature mixing of miscible LCLs into the downstream CC by using air as an isolation phase. (c) Multiphase domino valve (MDV) with an HCL upstream of the LCL reservoir. The MDV allows LCL reservoirs to be inserted into the MCR. (d) An integrated micromixer enables LNP fabrication by mixing lipids in an LCL and RNA in an HCL, which are simultaneously released into the mixer. Gray indicates empty microchannels, and color depth reflects microchannel depth. The HCL is shown in blue, the miscible LCL in orange, and the immiscible LCL in red.

### P2PV for LCL valving

The P2PV operates analogously to a conventional SV, stopping a liquid that fills a reservoir and preventing it from flowing uncontrollably into the main channel. A P2PV stops an LCL at the reservoir outlet using a hydraulic barrier, thereby confining a predefined volume of LCL within the reservoir. The filling sequence of an LCL reservoir with a P2PV differs from that of an HCL reservoir with conventional capillary valves. In a conventional HCL reservoir, an aqueous HCL is loaded directly into the HCL reservoir, spontaneously fills the reservoir and then the functional connection, and stops at the SV located between the functional connection and the main channel. Subsequent filling of the main channel with an aqueous HCL establishes hydraulic connectivity of the reservoir with a paper pump, enabling CC operation^22^. In contrast, LCL reservoir filling begins with prefilling the main channel with an aqueous HCL. The HCL then enters the functional connection and stops at an oppositely oriented SV located between the functional connection and the reservoir, thereby forming the control HCL, Fig. 2a. Next, the LCL is loaded into the reservoir inlet. However, air trapped between the HCL and the LCL would prevent the LCL from advancing through the reservoir and contacting the control HCL. We thus added two air vents, one on each side of the reservoir outlet to allow the air to escape. Each vent comprises an open-topped chamber with a smaller cross-section than the reservoir, connected by a narrow, shallow channel. As the LCL fills the reservoir and contacts the control HCL, the LCL is stopped and establishes a hydraulic connection with the CC, Fig. 2a. Although the LCL still bursts into the air vents during reservoir loading, the overflowed LCL does not flow back into the reservoir during delivery thanks to the smaller cross-section yielding a higher capillary pressure than the reservoir. Excessive LCL loading can saturate the air vents and compromise P2PV function; this could be prevented by incorporating a liquid-volume metering function^29^, which was not implemented in this study.

**Figure 2.**
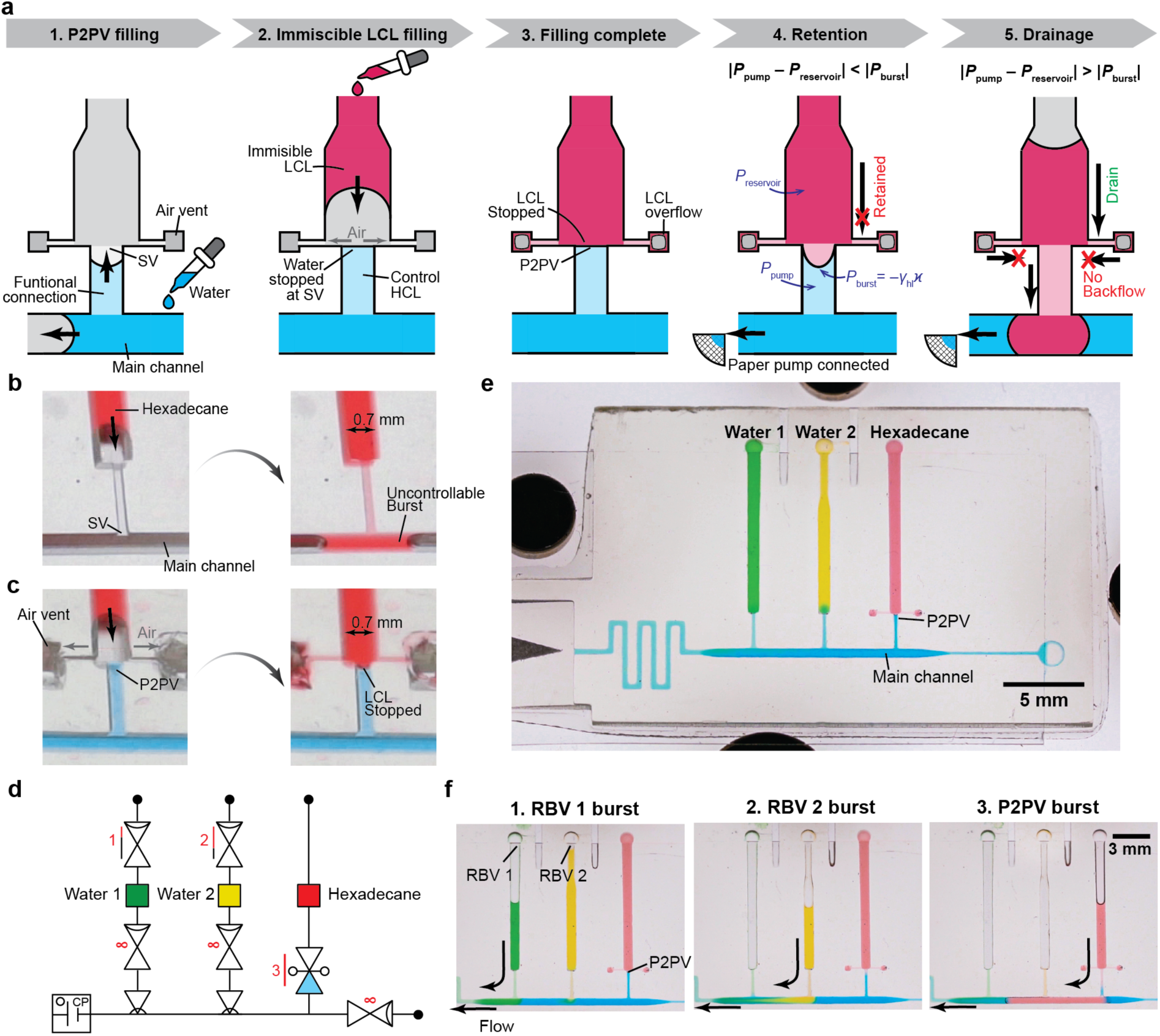
Sequential delivery of immiscible LCLs by a P2PV, Videos S1–S2. (a) P2PV workflow for the hydraulic control of an immiscible LCL. The P2PV uses the interface formed between the control HCL and the immiscible LCL to confine the LCL and encode a burst pressure for programmable sequential delivery. Gray indicates empty microchannels, and color depth reflects microchannel depth. (b) Failure of the SV in hexadecane confinement. (c) Successful confinement of hexadecane by the P2PV. (d) Circuit diagram and (e) corresponding 3D-printed CC used for the sequential delivery of Water 1 (green), Water 2 (yellow) and hexadecane (red) with volumes of 5, 3 and 5 µL, respectively. (f) Sequential delivery of the three liquids.

Programmable liquid delivery of LCLs requires different valving strategies depending on whether the LCL is immiscible or miscible with the control HCL of the P2PV. An immiscible LCL does not mix with the aqueous HCL and thus forms an interface at the functional connection. Analogous to an RBV, which uses the capillary pressure generated by the surface tension of the liquid-air interface, the P2PV uses the interfacial tension of the LCL-HCL interface at the functional connection for programmable retention and delivery. Upon bursting of the P2PV, the control HCL is released, entraining the LCL into the main channel. In contrast, a miscible LCL mixes with the control HCL and does not form an interface; consequently, the P2PV cannot retain miscible LCLs due to the negligible capillary pressure and the LCL can be mixed into the main channel. Programmable delivery of miscible LCLs is therefore achieved by adding an HPR downstream of the P2PV (Fig. 1b).

### Programmable sequential delivery of immiscible LCL by a P2PV

The process of reservoir filling and pre-programmed sequential delivery of an immiscible LCL is shown in Fig. 2a. An aqueous HCL first fills the main channel and then forms a control HCL in the functional connection. An immiscible LCL is loaded into the reservoir inlet and fills the reservoir while displacing air through the air vents. Upon contacting the control HCL, the LCL stops and a burst pressure arises as *P*_burst_ = –*γ*_hl_*κ* where *γ*_hl_ is the interfacial tension at the HCL-LCL interface, and *κ* is the interfacial curvature governed by the cross-section of the channel and the receding contact angle. Upon connection of the paper pump, the LCL is retained in the reservoir while |*P*_pump_ – *P*_reservoir_ | < |*P*_burst_|. When the burst threshold is reached, |*P*_pump_ – *P*_reservoir_ | > |*P*_burst_|, the control HCL bursts and then the LCL is released into the main channel, Fig. 2a.

We demonstrate the handling of hexadecane, an immiscible LCL, in CCs with P2PVs. Compared to water with a functional contact angle, *θ*_water_ = 46°, hexadecane (red) exhibited a substantially lower contact angle, *θ*_hexadecane_ = 22° (Fig. S1) below the functional range and was not retained by a conventional SV and after a short time, burst, and spread uncontrollably into the main channel, Fig. 2b. In contrast, a P2PV successfully confined hexadecane within the reservoir, Fig. 2c. The systematic failure of SVs and concomitant success of P2PVs are illustrated using a 3D-printed CC with ten interconnected reservoirs, five with an SV and five with a P2PV, Fig. S2 and Video S1.

We next demonstrate the coordinated sequential delivery of water and hexadecane within a single CC. Water reservoirs are controlled by conventional capillary valves including SVs, RBVs, and CRVs, whereas a hexadecane reservoir is controlled by a P2PV. The circuit diagram in Fig. 2d and the equivalent 3D-printed CC in Fig. 2e illustrate and implement this strategy. Water reservoirs 1 and 2 have an RBV with burst pressures at scales 1 and 2, respectively, while the hexadecane reservoir is controlled by a P2PV with a burst pressure at scale 3, enabling the sequential delivery of Water 1 – Water 2 – Hexadecane, Fig. 2f. Detailed operation including liquid loading and sequential delivery is shown in Video S2.

### Valving of miscible LCL while avoiding pre-mixing using P2PV-HPR

P2PVs can confine miscible LCLs, but upon contact between the HCL and LCL, mixing by diffusion and by surface-tension-driven Marangoni flow cause the LCL to reach the main channel via the functional connection and mix with the liquid therein rapidly, even in the absence of flow. This rapid mixing was visualized by using a fluorescent dye in ethanol and observing the fluorescence upon contact between the ethanol and the control HCL in the functional connection at *t* = 0, Fig. 3a. After as little as 1 s, fluorescence is already visible in the main channel, and continued to propagate. For many applications, this premature mixing could lead to unwanted interaction between solutions in different reservoirs and the main channel.

**Figure 3.**
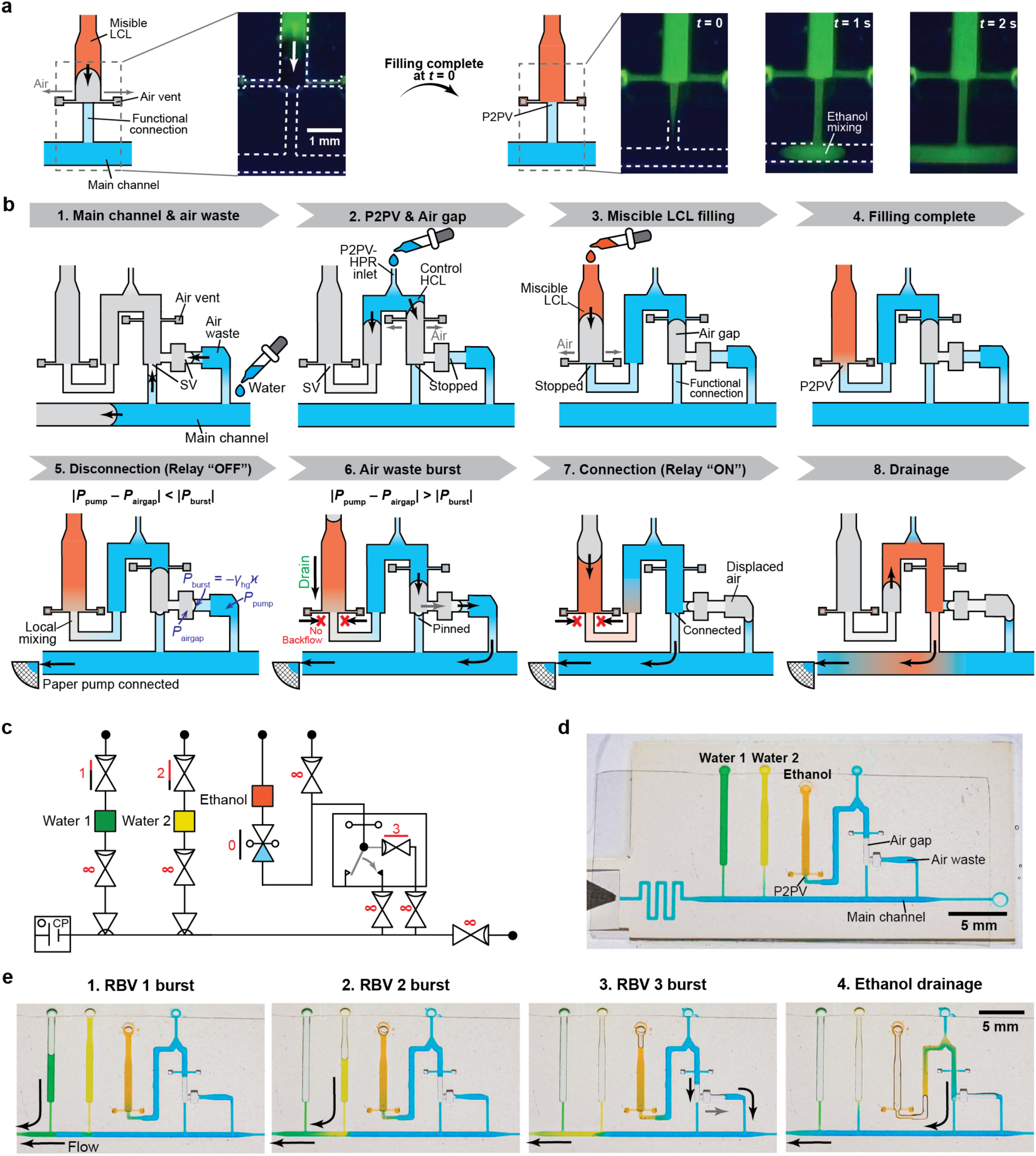
Miscible LCL valving using a P2PV-HPR, Video S3. (a) Ethanol loading into a reservoir connected with the main channel through a P2PV directly formed in the functional connection. Once the advancing ethanol is stopped by the P2PV at *t* = 0, ethanol mixes into the main channel by diffusion and Marangoni flow, with the mixing front advancing ∼1 mm within 1 s. Ethanol is shown in orange in the schematics and green in the fluorescence images. Dotted lines in the fluorescence images indicate channel walls. (b) Workflow of miscible LCL valving using a P2PV-HPR. The air gap initially disconnects the LCL reservoir from the main channel, enabling loading and retention without premixing. An RBV formed in the air waste defines the burst threshold for programmed delivery. Upon reaching the burst condition, the RBV bursts, the air waste accommodates the displaced air gap and LCL reservoir is connected to the main channel for delivery. Gray indicates empty microchannels, and color depth reflects microchannel depth. (c,d) Coordinated sequential delivery of two water reservoirs (green and yellow) and one ethanol reservoir (orange) using the P2PV-HPR, shown as (c) a circuit diagram and (d) the corresponding 3D-printed CC. The reservoirs contain Water 1 (green), Water 2 (yellow) and ethanol (orange) with volumes of 5, 3 and 2.5 µL, respectively. (e) Sequential delivery of the three liquids.

To prevent premature mixing, we introduce the P2PV-HPR by integrating a HPR consisting of an air gap and a water-filled air waste (Fig. 1b) downstream of the P2PV. The air gap isolates the LCL reservoir from the main channel and restricts premixing within the HPR. At time of flow controlled by an RBV retaining an HCL in the air waste, the air is directed into the air waste and hydraulic connection of the miscible LCL to the main channel is realized only during the delivery. Figure 3b illustrates loading and delivery process of the P2PV-HPR for miscible LCL: (1) Water is first introduced into the main channel, and as described previously, fills the functional connection as well as the air waste up to the SVs. (2) The control HCL is loaded into the P2PV-HPR inlet, flowing both upstream to the SV to prime the P2PV, and downstream to trap air thus forming an air gap in the HPR. (3) The miscible LCL is then loaded into the reservoir and stops at the P2PV, (4) where it contacts the water and premixes with it. However, mixing is restricted to the water plug and does not spill into the main channel because the air gap disconnects the relay (“OFF” state). In our experiments, no visible ethanol mixing in the main channel was observed for at least 5 min (data not shown). (5) Upon paper-pump actuation of the CC, and (6) upon exceeding the burst pressure *|P*_pump_ *– P*_airgap_*| > |P*_burst_| of the air waste (*P*_burst_ = –*γ*_hg_*κ* where *γ*_hg_ is the surface tension at the HCL-gas interface), the trapped air is hydro-pneumatically driven into the air waste until (7) the control HCL merges with the liquid in the functional connection, switching the relay to the “ON” state. The LCL is now connected hydraulically to the main channel and the paper pump, and (8) flows until the reservoir is emptied.

Next, we integrated the P2PV-HPR into a CC for the coordinated sequential delivery of water and (miscible) ethanol as illustrated schematically in Fig. 3c and implemented in the 3D-printed chip in Fig. 3d. The delivery sequence, Water 1–Water 2–Ethanol was as pre-programmed using progressively increasing RBV burst-pressure scales from 1–3, where scale 3 applies to the air waste coupled to the ethanol reservoir, Fig. 3e. Detailed operation is shown in Video S3.

### Multiphase MCR

The MCR structurally encodes the sequential delivery of liquids from *N* reservoirs by imposing the condition that reservoir *n* will be drained only after reservoir *n–*1 has been drained, realizing a cascading liquid delivery in CCs, Fig. 4a^23^. Conventionally, capillary domino valves (CDVs) gate the self-propagation of MCRs, and have an air link between adjacent reservoirs, bookended by a transmitter valve (a combined SV and RBV) at the outlet of reservoir *n*–1 and a receiver valve (a combined SV and RBV) at the inlet of reservoir *n*. However, the SV again restricts the functionality of MCR to HCLs because the CDVs comprise SVs that are not operational with LCLs. Here, we extend MCR operation to multiphase liquid handling by integrating a P2PV-HPR into the CDV to make it an MDV, Fig. 4b. The water located upstream of the LCL reservoir defines a receiver valve for LCL reservoir *n*, which, together with a transmitter valve of the preceding reservoir *n*–1, forms an MDV. The MDV enables the incorporation of an LCL at any position *n* within an MCR and supports the processing of both immiscible and miscible LCLs, Fig. 4b.

**Figure 4.**
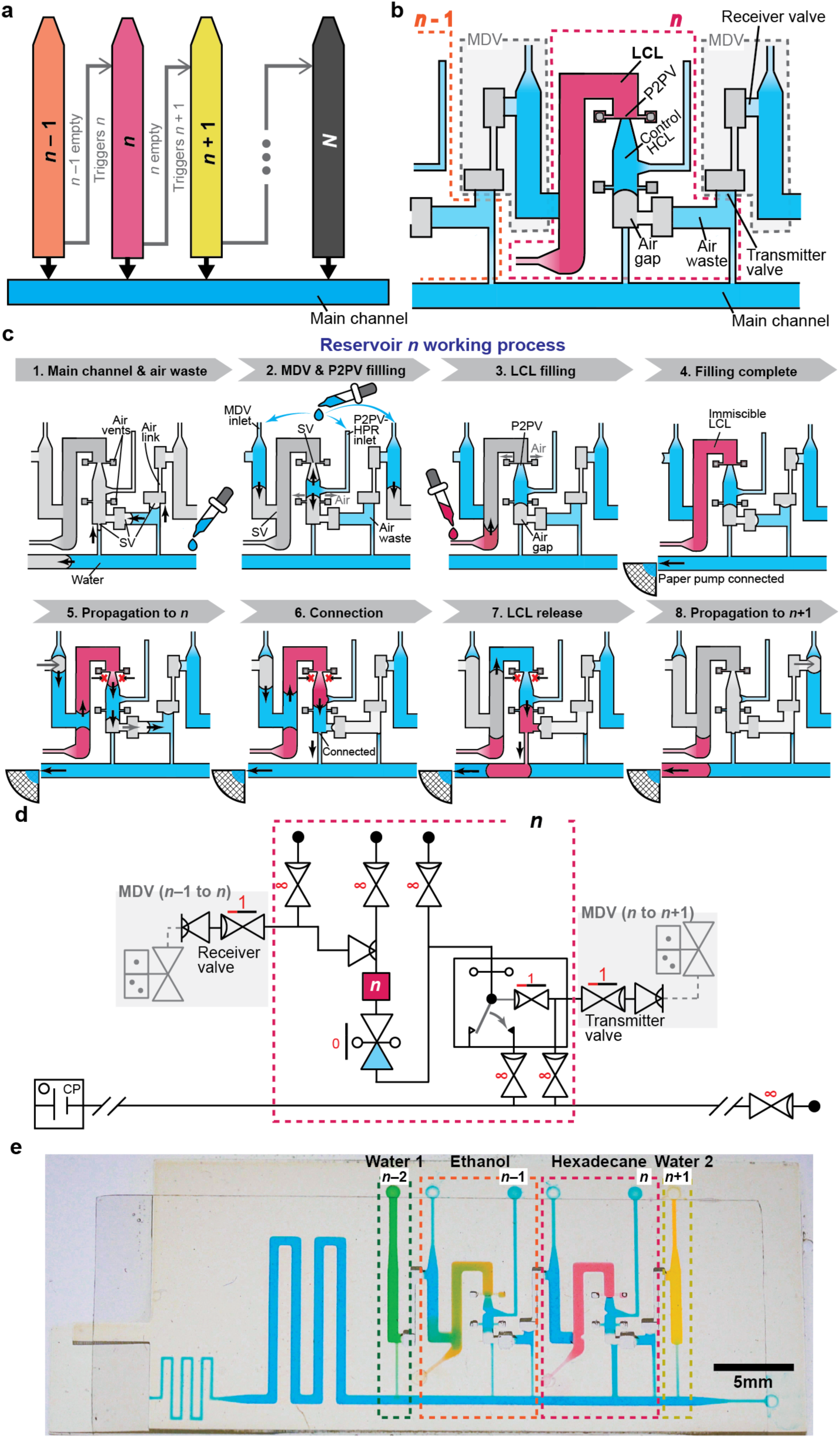
Multiphase microfluidic chain reaction (MCR), Video S4. (a) Concept of multiphase MCR for serial release of liquids, in which drainage of one reservoir triggers release of the following reservoir through multiphase domino valves (MDVs). (b) Schematic structure of a multiphase MCR. An LCL reservoir at event *n*, is integrated into the MCR between reservoirs *n*–1 and *n*+1 through an MDV and a downstream P2PV-HPR architecture. Both immiscible and miscible LCLs are controlled by the same architecture. (c) Working process of the MCR at reservoir *n* containing an immiscible LCL (red). Connection of air to the receiver valve triggers connection of the LCL reservoir to the main channel by the HPR, initiating LCL delivery. Subsequent drainage of the HCL completes event *n* and propagates the air connection to reservoir *n*+1. Gray indicates empty microchannels, and color depth reflects microchannel depth. (d,e) Multiphase MCR of water (green and yellow), ethanol (orange), hexadecane (red), with each reservoir loaded with 2.5 µL, along with 1.4 µL MDV and 0.5 µL control HCLs for each LCL reservoir, shown as (d) a circuit diagram and (e) the 3D-printed multiphase MCR CC.

Figure 4c illustrates the loading and operation of a reservoir *n* containing an LCL within a multiphase MCR: (1) First, the HCL is added to the main channel, filling the functional connection and the air waste, followed by (2) the addition of the HCL into the P2PV-HPR and MDV inlets, and (3) loading of the LCL, which primes the reservoir. (4) Upon completing the filling process, (5) the CC is actuated by connecting a paper pump and complete drainage of reservoir *n*–1, leads to connection of the air pathway to the receiver valve of reservoir *n*. (6) The HPR switches from “OFF” to “ON”, hydraulically connecting the LCL to the main channel, and (7) the LCL is drained together with the MDV and control HCLs into the main channel until (8) all liquid is drained, thus opening an air pathway to reservoir *n+*1. A circuit diagram of an LCL reservoir *n* within an MCR and the corresponding 3D-printed CC are shown Fig. 4d,e. Detailed operation is shown in Fig. S3 and Video S4. The multiphase MCR sequentially delivers water (green and yellow), ethanol (orange) and hexadecane (red), demonstrating propagation across all combinations of phase transitions: HCL-to-LCL, LCL-to-LCL, and LCL-to-HCL. These results establish the multiphase MCR as a platform capable of sequential delivery of arbitrary combinations of HCLs and LCLs.

### Multiphase MCR CC for autonomous on-chip lipid nanoparticle fabrication

We illustrate the potential of multiphase MCR CCs for autonomous, structurally pre-programmed LNP fabrication. LNPs are typically made by controlled volumetric mixing of three parts of an aqueous solution (e.g. acidic citrate buffer) containing the RNA cargo and one part of a solvent (e.g. ethanol) containing lipids, which induces the rapid self-assembly of LNPs encapsulating the RNA. To date, LNP fabrication commonly relies on automated mixing systems spanning microliter-to-milliliter working volumes for formulation development and substantially larger volumes for preclinical and commercial production. At the research scale, minimizing the working volume is particularly advantageous for formulation screening because RNA cargo and lipid reagents are often costly. Conventional microfluidic systems, however, are dependent on peripherals including active pumps and controllers and computers and can incur reagent losses of tens to hundreds of microliters due to dead volume and the establishment of stable flow, making small-volume formulation particularly inefficient^4–6^, Fig. 5a. Further, RNA samples often require pre-dilution prior to mixing, which, because of the complexity of conventional microfluidic systems, is often performed offline.

**Figure 5.**
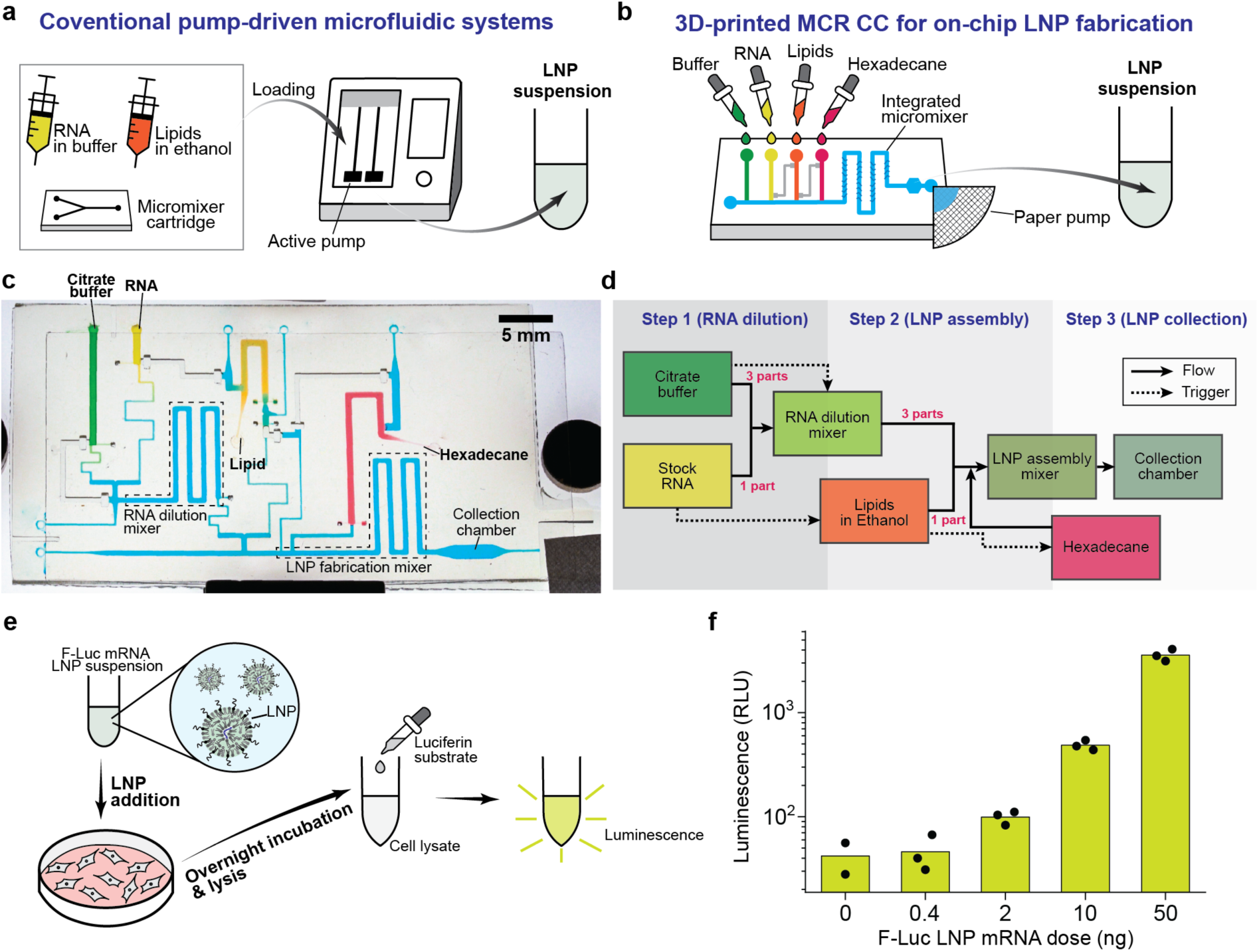
Autonomous on-chip lipid nanoparticle (LNP) fabrication using a multiphase MCR CC, Video S5. (a) Conventional pump-driven microfluidic systems. RNA and liquid solutions are loaded with a micromixer cartridge and flow is driven by active pumps at actively controlled flow rates. (b) Autonomous LNP fabrication in a multiphase MCR CC. The aqueous RNA solution (3.6 µL), buffers (10.8 µL), lipids in ethanol (5.2 µL) and hexadecane (12.5 µL) are loaded on-chip, and upon paper-pump connection, LNPs are autonomously fabricated and collected. (c) 3D-printed multiphase MCR CC for on-chip LNP fabrication. (d) Flowchart for LNP fabrication using the multiphase MCR CC. (e) mRNA-LNP cell transfection workflow. LNPs encapsulating firefly luciferase (F-Luc) mRNA were added to cells cultured in a well plate. After overnight incubation and cell lysis, luciferin was added to the cell lysate, and the luminescence generated by F-Luc activity was measured. (f) Luminescence signals from cell lysates following transfection with F-Luc mRNA-LNPs at varying total mRNA inputs; the corresponding encapsulated doses are determined from the measured encapsulation efficiency of 61 ± 2%. Bars indicate the means and points show individual replicates (*n* = 3, except *n* = 2 at 0 mRNA).

Here, we introduce a 3D-printed multiphase MCR CC that enables simplified LNP fabrication without need for peripherals, using a total formulation volume of only ∼20 µL. Buffer, stock RNA solution, lipid solution and hexadecane are loaded into their respective reservoirs. The device integrates micromixers for RNA dilution and LNP fabrication, both incorporating staggered herringbone structures to promote chaotic mixing, which are widely used in microfluidics because of their relative ease of fabrication. Upon actuation by paper-pump connection, the MCR autonomously dilutes the stock RNA, mixes it with the lipids in ethanol and delivers the assembled LNPs to an on-chip collection chamber for retrieval, Fig. 5b–d.

The CC comprises two aqueous reservoirs containing 3.6 µL of stock RNA solution in water (yellow) and 10.8 µL of citrate buffer, pH 4 (green), respectively, and two LCL reservoirs containing 5.2 µL of lipid solution in ethanol (orange) and 12.5 µL of hexadecane (red), respectively, Fig. 5c. The main channel, RNA dilution mixer, and LNP fabrication mixer are filled with the priming buffer (citrate buffer). As summarized in the flowchart in Fig. 5d, the stock RNA solution and the citrate buffer are first released simultaneously into the RNA dilution mixer, diluting the RNA 1:4 to prepare the 14.4 µL of working RNA solution at pH 4. Complete drainage of the RNA and buffer reservoirs subsequently triggers the simultaneous release of the lipid reservoir and the RNA dilution mixer, respectively, into the LNP fabrication mixer. At the beginning of this step, small leading volumes of the priming buffer in the RNA dilution mixer and the control HCL for the lipid reservoir are discharged while these volumes were accounted for in the device design. The working RNA and lipid solutions then enter the LNP fabrication mixer at a 3:1 flow rate ratio (RNA:lipid), initiating LNP fabrication. Complete drainage of the lipid reservoir along with its aqueous MDV HCL, triggers the release of hexadecane into the LNP fabrication mixer. The hexadecane, as an immiscible phase, then displaces the LNP suspension into the 6 µL collection chamber. The chamber therefore retains only the final 6 µL fraction, while the preceding volume including the fraction most affected by the priming liquids, is discarded into the paper pump. The autonomous on-chip operation of LNP synthesis is demonstrated in Fig. S4 and Video S5. The LNPs were retrieved with a needle used to pierce the tape cover of the chamber and a pipette to aspirate the LNP solution from the collection chamber.

The desired flow rates and ratios were obtained by first analytically predicting the flow rates based on the viscosity of each solution and the channel geometry, 3D printing chips accordingly, and experimentally verifying the predictions by measuring flow times, followed by iterative fine-tuning of the fluidic resistances of the different channels in the network (Fig. S4).

The herringbone mixer can act as an unintended SV to aqueous HCL that is used to prime the CC, thereby interrupting capillary filling. The abrupt edges of grooves can pin the advancing meniscus, or can trap bubbles within the recesses, thus compromising the mixing and overall chip functionality. We addressed these challenges by modifying the cross-sectional profile of the grooves along the flow direction. Each groove was designed to widen gradually on the upstream side to promote capillary filling of the recess, while the downstream side retained an abrupt, right-angled edge to perturb the streamlines and thus promote chaotic mixing. The mixer was characterized and mixing of ∼80% was achieved at ∼20 s after the liquids entered the micromixer at a flow speed of ∼0.67 mm/s, which was deemed adequate for our application (See Fig. S5 for further details).

The RNA-LNPs made using the CC exhibited a mean diameter of 157 ± 75 nm, a zeta potential of –6.8 ± 0.8 mV, and an RNA encapsulation efficiency of 61 ± 2%. Although these results confirm successful RNA-LNP fabrication, the relatively large particle size, broad size distribution, and marginally lower encapsulation efficiency indicate that the formulation performance requires further optimization. Possible contributing factors include: (i) the comparatively slow mixing in the CC resulting from its low flow rate, with a residence time of ∼70 s and ∼80% mixing achieved after ∼20 s, which is substantially longer than conventional LNP mixers with the millisecond timescale^30^, and (ii) dilution of the RNA and lipid solutions by the priming buffer prefilled in the mixers, which may reduce the effective reagent concentrations. Nevertheless, the CC performed the complete fabrication process using a total working volume of only ∼20 µL, demonstrating that autonomous LNP fabrication can be implemented at small scale without external pumps or controllers. To assess biological activity of the LNPs fabricated using the CC, HEK293T cells were transfected with LNPs containing firefly luciferase messenger RNA (F-Luc mRNA), and luciferase expression was measured using a luminescence assay after overnight incubation (Fig. 5e). Increasing the total mRNA dose produced a corresponding increase in luminescence, confirming the dose-dependent intracellular delivery and translation of the encapsulated mRNA, Fig. 5f. These results establish the multiphase MCR CC as a proof-of-concept for the autonomous implementation of complex multiphase nanomedicine workflows.

## Conclusion

We introduced a library of multiphase capillaric components and architectures for coordinated multiphase liquid handling in CCs. P2PVs use a control HCL to shift capillary control from gas-liquid interfaces, as employed in conventional capillary valves including SVs, RBVs, and CRVs, to liquid-liquid interfaces, enabling LCLs to be hydraulically confined and released in CCs. P2PVs enabled the pre-programmed handling of immiscible LCLs and P2PV-HPR architectures extended this capability to miscible LCLs while preventing premature mixing into the downstream CC before actuation. We further introduced MDVs to integrate LCL reservoirs into the MCR with P2PV-HPR architectures, extending the MCR from an aqueous system to a multiphase system capable of propagating sequential and simultaneous liquid release across HCL-LCL, LCL-LCL, and LCL-HCL transitions. We demonstrated an LNP fabrication MCR CC that executes stock RNA dilution, LNP fabrication by mixing RNA in buffer and lipids in ethanol, and product collection by oil, demonstrating that multiphase CCs can reproduce biochemical manufacturing workflows. More broadly, this work expands the capillaric toolbox beyond predominantly aqueous operation by adding multiphase components and circuit architectures to the existing repertoire of valves and advanced circuit elements. By enabling autonomous control and coordination across miscible and immiscible LCL phases, this work broadens the design space of capillaric systems and the range of multistep fluidic workflows that can be implemented in 3D-printed CCs.

## Materials and methods

### Materials

#### Capillaric circuit fabrication and fluidics demonstration

ABS-like resin Pro 2, Clear (Cat. #SABP2CL-10-4, Anycubic, Shenzen, China), Isopropyl alcohol (IPA) (Cat. BDH1133, VWR, Radnor, PA, USA), Hexadecane (Cat. #H6703, lot #SHBP8192, Sigma-Aldrich, Oakville, ON, Canada), Ethanol anhydrous (Cat. #P016EAAN, Greenfield Global, Brampton, ON, Canada), 2,3-butanediol (Cat. #B84094, lot #BCBG4025V, Sigma-Aldrich, Oakville, ON, Canada), Oil red O (Cat. #O9755, lot #SHBQ7480, Sigma-Aldrich, Oakville, ON, Canada), Sudan I (Cat. #103624, lot #BCBZ5090, Sigma-Aldrich, Oakville, ON, Canada), Coumarin 153 (Cat. #546186, lot # MKCW3868, Sigma-Aldrich, Oakville, ON, Canada), Fluorescein sodium salt (Cat. #46960, lot #2082530, Sigma-Aldrich, Oakville, ON, Canada).

#### LNP fabrication, characterization and transfection

SM102 (Cat. #33474, lot #0650585-9, Cayman Chemical, Ann Arbor, MI, USA), Cholesterol (Cat. # C8667-5G, lot #SLCG0969, Sigma-Aldrich, Oakville, ON, Canada), DSPC (Cat. #850365P-1g, lot #850365P-1G-X-178, Avanti Polar Lipids Inc., Alabaster, AL, USA), DMG-PEG2000 (Cat. # 880151P-1g, lot # 880151P-1G-A-025, Avanti Polar Lipids Inc., Alabaster, AL, USA), Ethanol anhydrous, F-Luc mRNA (Cap1, m1Ψ) (Cat. #RP-A00023-0.2, lot # U5004695G0, GenScript, Piscataway, NJ, USA), Sodium citrate, 0.5M buffer soln., pH 5.0 (Cat. #J62918.AP, lot #Y29K526, Thermo Fisher Scientific, Waltham, MA, USA), Nuclease-free water (Cat. #W4502, Lot #0000377069, Sigma-Aldrich, Oakville, ON, Canada), Triton X-100 (Cat. #BP151-500, lot #051126, Thermo Fisher Scientific, Waltham, MA, USA), Quant-iT RiboGreen RNA Assay Kit (Cat. #R11490, lot #2600134, Thermo Fisher Scientific, Waltham, MA, USA), Luciferase Assay System (Cat. #E4030, lot #0000690354, Promega, Madison, WI, USA).

### Methods

#### CC design and fabrication

CC designs were prepared using Fusion 360 (Autodesk, San Francisco, CA, United States), exported as *.stl* files and fabricated using an LCD 3D printer^27^ (Elegoo Mars 4 Ultra, Elegoo, Shenzhen, China) with a ABS-like clear resin. Printed CCs were then rinsed with IPA, and post-cured for 1 min under UV illumination using a UV chamber (Professional CureZone, Creative Cadworks, Toronto, ON, Canada). For surface treatment, CCs were then plasma-treated for 10 s at 30% power (PE50 plasma chamber, Plasma Etch, Carson City, NV, USA) and sealed with a pressure-sensitive adhesive tape (9795R microfluidic tape, 3M, Maplewood, MN, USA) which covered the open microchannels to form enclosed channels.

#### Liquid contact angle measurement

Contact angles were measured by capturing side-view images of droplets (*n* = 5) on plasma-treated CC surfaces using a digital camera (Panasonic Lumix DMC-GH3K, Panasonic, Osaka, Japan) equipped with a macro lens (M.Zuiko Digital ED 60 mm F2.8 Macro, Olympus, Tokyo, Japan). The images were analyzed using the contact angle extension in ImageJ.

#### Fluidic operation and demonstration

Fluidic demonstrations were performed using hexadecane and ethanol, representing water-immiscible and miscible LSLs, respectively as well as water. Ethanol was supplemented with 10% (v/v) 2,3-butanediol to suppress evaporation during fluidic demonstrations. For flow visualization, water, hexadecane and ethanol were dyed with 2% (v/v) food dye, 0.1 mg mL^-1^ oil red O and 0.1 mg mL^-1^ Sudan I, respectively. For fluorescence imaging, 0.1 mg ml^-1^ coumarin 153 in ethanol and a 100 µM fluorescein solution in water were used to visualize ethanol mixing into the main channel and mixing performance of the micromixer, respectively.

For CC operation, liquids were pipetted into the inlets of the CC, and once all reservoirs and the main channel were filled, the outlet was connected to a paper pump (Whatman filter paper grades 5 and 1, cat. #1005-125 and 1001-150, lot #10310096 and 16958998, GE Healthcare, Chicago, IL, USA). Videos and brightfield images were recorded using a digital camera (Sony α7R III, Sony, Tokyo, Japan) equipped with a macro lens (Sony FE 90 mm F2.8 Macro G OSS Lens, Sony, Tokyo, Japan). Fluorescence images were acquired using an inverted fluorescence microscope (Ti2, Nikon, Tokyo, Japan) with a 10× objective and an sCMOS camera (Prime 95B, Teledyne Photometrics, Tucson, AZ, USA) at an exposure time of 50 ms. The mixing index was calculated according to the conventional definition:

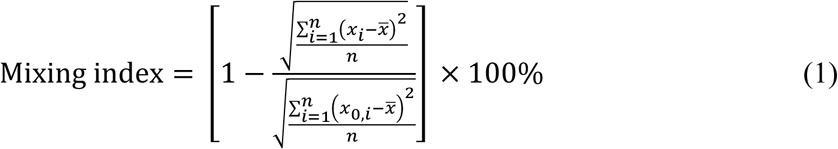

where *x_i_* is the intensity of pixel *i*, *x*_0,*i*_ is the intensity of pixel *i* at the entrance of the mixer, *x̄* is the average intensity, and *n* denotes the number of pixels sampled.

#### Flow rate analysis

Flow rates of the LNP CC were quantified from backlit video recordings by analyzing the time-dependent light transmission through user-defined channel regions. Variations in transmitted light intensity caused by changes in fluid occupancy within the optical path were used as an optical proxy for the liquid presence in the channel over time. For each region of interest, pixel intensities were extracted to generate intensity-time traces. A background-drift correction was applied to compensate for fluctuations in illumination and camera exposure during recording. The corrected intensity signals were subsequently normalized and converted to channel occupancy and calibrated liquid volume. Flow rates were calculated from the time-dependent volume profiles, generating synchronized volume and flow-rate traces for quantitative analysis.

#### LNP fabrication and analysis

For LNP fabrication, a 10 mM lipid solution composed of SM102, Cholesterol, DSPC, DMG-PEG2000 at a molar ratio of 50:38.5:10:1.5 was prepared in ethanol. An F-Luc mRNA solution was prepared at 360 µg/ml in nuclease-free water and 67 mM sodium citrate buffer was prepared at pH 4, so that after on-chip dilution, the RNA and lipid concentrations yielded a molar ratio of protonatable amine groups (N) on the ionizable lipid to phosphate groups (P), also known as N/P ratio, of 6:1.

For operation, 3.6 µL of stock RNA solution, 10.8 µL of 67 mM citrate buffer, 5.2 µL of lipid solution and 12.5 µL of hexadecane were loaded in their respective reservoirs while the main channel, RNA dilution mixer, and LNP fabrication mixer were primed with 50 mM citrate buffer. Upon paper-pump connection, the CC sequentially and simultaneously released liquids to implement RNA dilution, LNP fabrication and product collection. The RNA dilution mixer had a total volume of 15 µL, while the simultaneous release of 3.6 µL of stock RNA solution and 10.8 µL of citrate buffer generated 14.4 µL of working RNA solution, leaving 0.6 µL of priming buffer ahead of the solution in the mixer. The lipid reservoir similarly contained a 0.5 µL control HCL ahead of the lipid solution. Upon simultaneous triggering of release of the RNA dilution mixer and lipid reservoir, the 0.6 µL of priming buffer and 0.5 µL control HCL were discharged first, followed by the working RNA and lipid solutions. After complete drainage of the lipid reservoir, 12.5 µL of hexadecane was released and the LNP product was displaced into the collection chamber that can hold the final volume of 6 µL LNP suspension. To retrieve the fabricated LNPs, the sealing tape covering the collection chamber was pierced with a needle, and then the LNP suspension was collected using a pipette. LNP suspensions from five CCs were pooled, diluted 50-fold in PBS at pH 7.4, and filtered using an Amicon® Ultra Centrifugal Filter, 100 kDa MWCO (Cat. #UFC5100, lot #0000280113, Sigma-Aldrich, Oakville, ON, Canada) for buffer exchange.

LNP size distribution and zeta potential were measured using a ZetaView x20 instrument (Particle Metrix, Inning am Ammersee, Germany). RNA encapsulation efficiency was determined using a standard RiboGreen assay. RNA concentrations of the LNP suspensions with and without addition of Triton X-100 to the final concentration of 0.5 % (v/v) followed by 15 min incubation at room temperature, were measured and considered as total mRNA concentration [mRNA]_total_ and free mRNA concentration [mRNA]_free_. RNA encapsulation efficiency was calculated as ([mRNA]_total_–[mRNA]_free_)/[mRNA]_total_.

For transfection studies, HEK293T cells cultured in a 96-well plate were treated with LNPs at different LNP doses and incubated overnight. The cells were subsequently lysed and luciferase activity of cell lysates was quantified using a luciferase assay kit and a plate reader (FLUOstar OPTIMA, BMG LABTECH, Ortenberg, Germany) according to the manufacturer’s instructions.

## Acknowledgement

We thank the Center for Applied Nanomedicine (CAN) at the RI-MUHC for access to the NTA system and support. This research was supported by The Natural Sciences and Engineering Research Council of Canada (NSERC) Discovery Grant RGPIN-2022-05171, Canada Research Chair in Bioengineering CRC-232159. G.K. acknowledges an FRQNT doctoral research scholarship and McGill BME recruitment and excellence awards. M.L.S acknowledges a Vanier Canada Graduate Scholarship and a McGill BME recruitment award.

## Conflict of Interest

The authors have no conflict of interest to declare.

## Author contributions

Conceptualization: G.K., A.N., D.J.

Methodology: G.K., A.N., D.J.

Investigation: G.K., L.A.H, M.L.S., Y.M.

Visualization: G.K., L.A.H, M.L.S., Y.M.

Funding acquisition: A.N., D.J.

Supervision: D.J.

Writing – original draft: G.K., A.N., D.J.

Writing – review & editing: G.K., L.A.H, M.L.S., Y.M., A.N., D.J.

## Data Availability

All datapoints are presented in the article and Supplementary Information. Source data will be available upon request. 3D design files will be uploaded to Thingiverse and Printables upon publication. (https://www.thingiverse.com/junckerlab/collections and https://www.printables.com/@JunckerLab_743461).

## Additional Information

Supplementary Information is available for this paper. Correspondence and requests should be addressed to

## Notes

### Competing Interest Statement

The authors have declared no competing interest.

