## Supplementary Information for "Multiphase capillarics for structurally pre-programmed sequential and parallel operations of high and low cohesion liquids"

### Supplementary figures

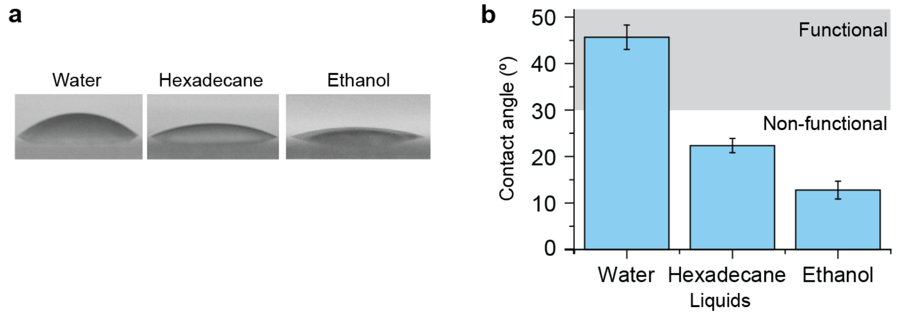


**Figure S1. Contact angles of water, hexadecane and ethanol.** (a) Side-view images of liquid droplets. (b) Measured contact angle of water, hexadecane and ethanol. The empirically observed functional contact angle range for capillary filling and valving, ~30–60º, is indicated by gray-shaded area.

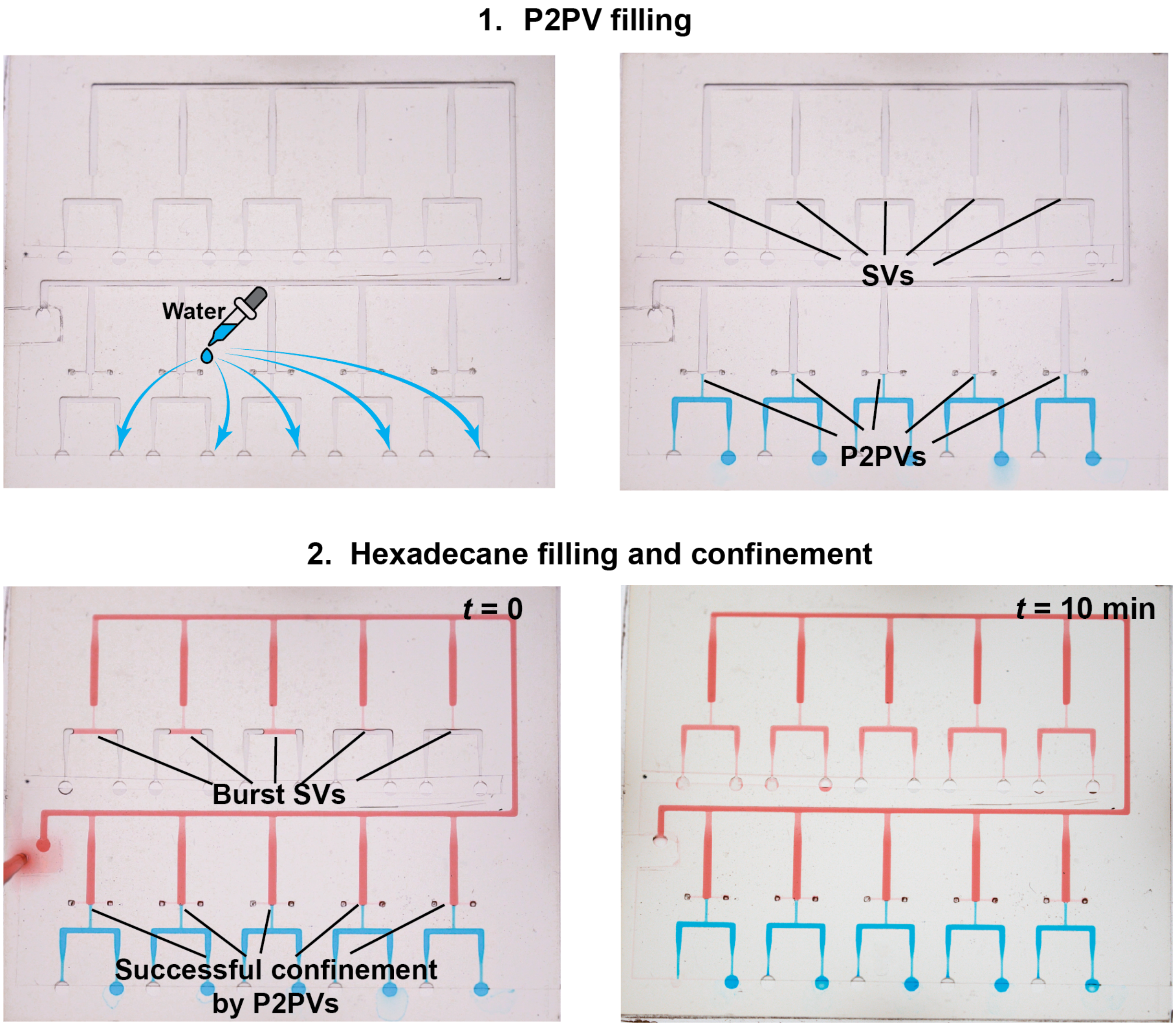


**Figure S2. Hexadecane confinement in serial reservoirs.** Five reservoirs with either an SV or a P2PV, respectively, are filled with hexadecane from a shared inlet. SVs fail to confine hexadecane and uncontrollably burst, while P2PVs successfully confine hexadecane, thus allowing progressive reservoir filling with pre-defined volumes. The volume of each reservoir is 2.5 µL.

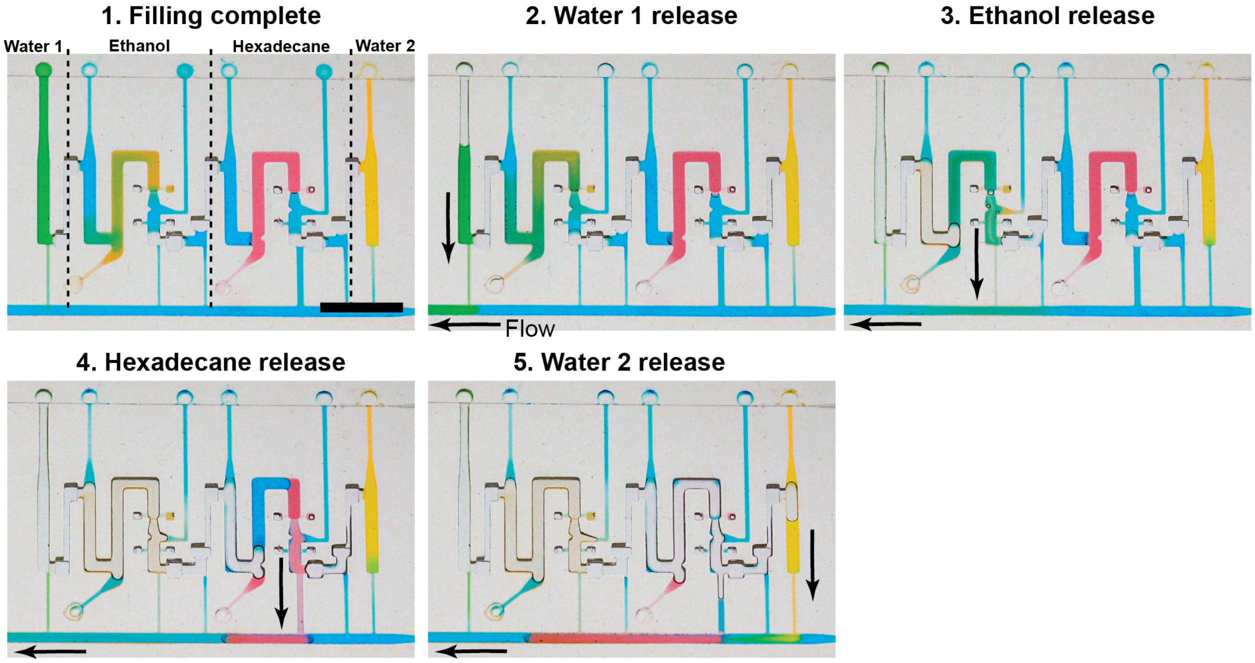


**Figure S3. Multiphase MCR for water, ethanol and hexadecane.** Liquids are each loaded at 2.5 µL along with 1.4 µL MDV and 0.5 µL control HCL for LCLs. Upon paper-pump connection, water, ethanol and hexadecane are sequentially released in pre-programmed sequence. Scale bar = 5 mm.

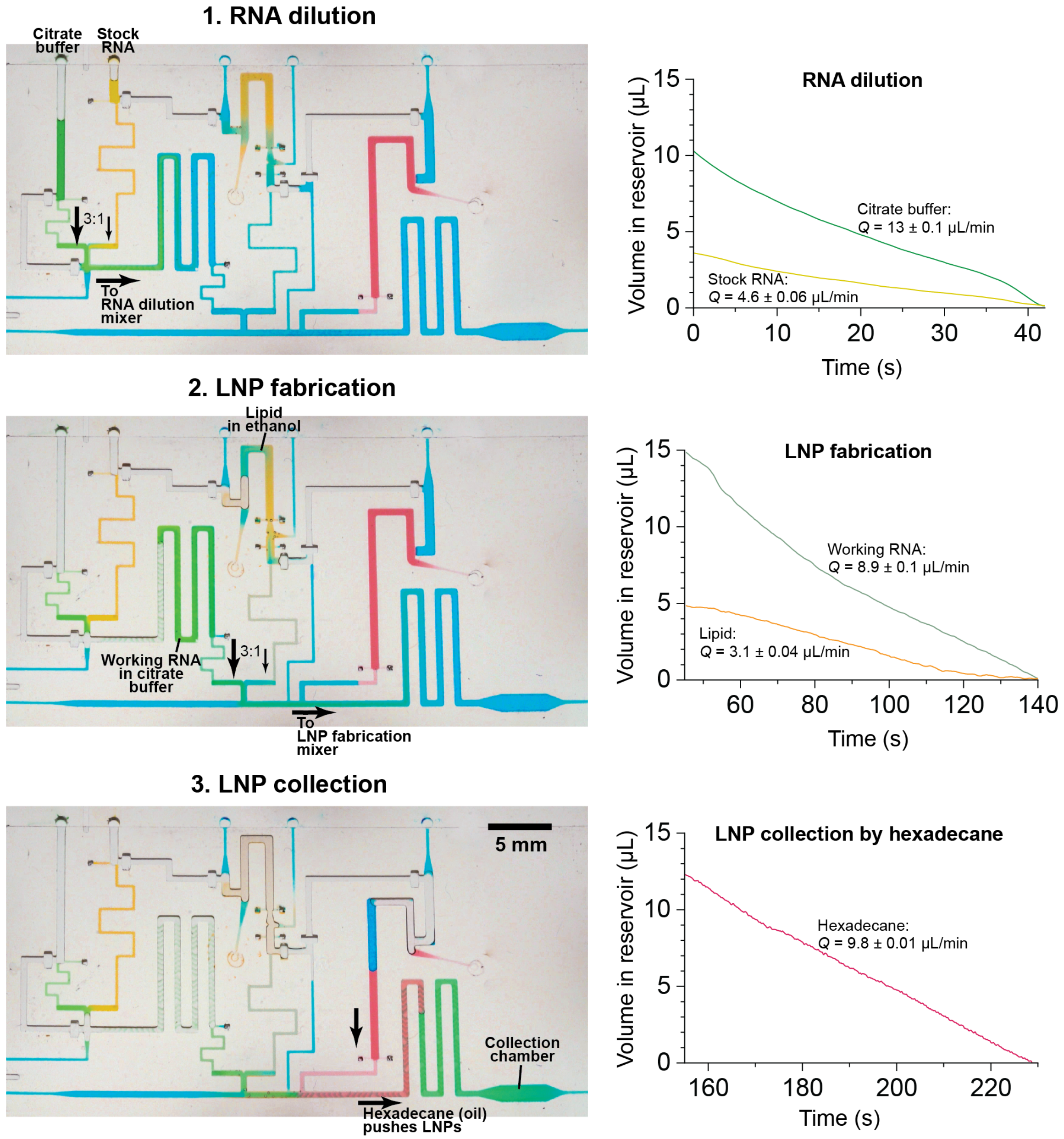


**Figure S4. MCR CC for LNP fabrication.** First, the 3.6 µL stock RNA solution is 1-in-4 diluted in 10.8 µL citrate buffer in the RNA dilution mixer, and the 14.4 µL working RNA solution is then mixed with the 5.2 µL lipid solution in ethanol in the LNP fabrication mixer at a flow-rate ratio of 3:1 (RNA:lipid) for RNA-LNP assembly. Finally, 12.5 µL hexadecane is released into the LNP fabrication mixer to displace LNPs into the 6 µL collection chamber, discarding first fraction of the LNP suspension. LNPs are then retrieved by piercing the sealing tape on the collection chamber.

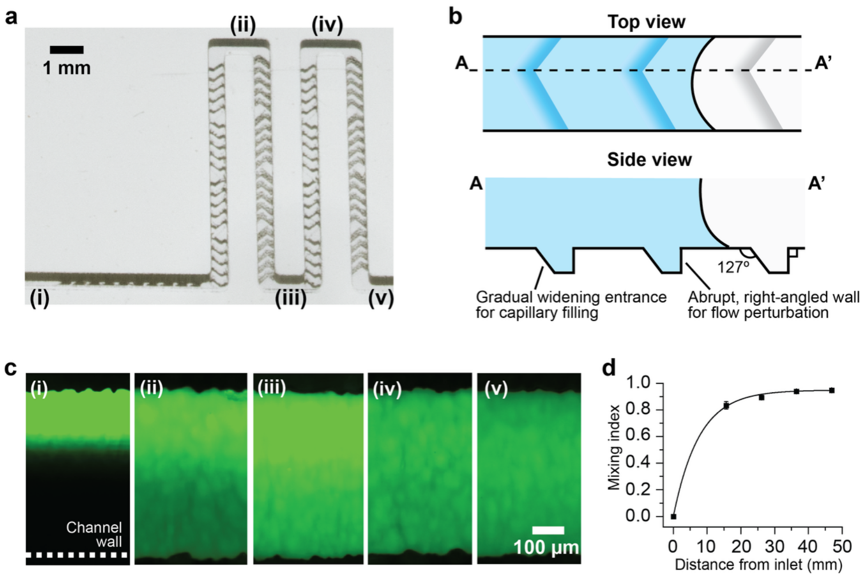


**Figure S5. Capillary staggered herringbone mixer.** (a) Serpentine micromixer with a staggered herringbone pattern on the channel bottom. (b) The gradually widening entrance allows smooth capillary filling of the recesses while the abrupt, right-angled wall promotes flow perturbation for chaotic advection. (c) Mixing of fluorescein solution and blank water at a 1:3 flow-rate ratio. Fluorescence images, acquired at the five locations indicated in **a**, show progressive micromixing along the serpentine channel. (d) Mixing index as a function of distance from the mixer inlet.

### Supplementary Videos

**Video S1. LCL confinement in interconnected reservoirs: P2PV versus SV.** Ten reservoirs, five incorporating an SV and five incorporating a P2PV, are interconnected in series and share a common inlet on a 3D-printed CC. The P2PVs are formed by introducing the control HCL into the functional connection. The HCL stops at the SV located at the junction between the functional connection and the reservoir. Hexadecane is loaded into the shared inlet and fills the reservoirs. In the reservoirs with an SV, hexadecane flows directly into the reservoir and the functional connection and encounters the SV at an abrupt widening into another channel. Due to the low cohesion, hexadecane bursts uncontrollably through the SV and spreads strongly. Hexadecane also fills the reservoirs with a P2PV by displacing the air through the air vents and upon contact, the P2PV successfully confines hexadecane within the reservoirs.

**Video S2. Coordinated handling of water and hexadecane as an immiscible LCL by the P2PV.** The 3D-printed CC implements the sequential delivery of three reservoirs including Water 1 (green), Water 2 (yellow) and Hexadecane (red) as an immiscible LCL. Water reservoirs are filled first, the main channel and then the functional connection are filled with the control HCL, forming a P2PV for the hexadecane reservoir and finally hexadecane is loaded. Upon connection of a paper pump, the liquids are sequentially released in the following order: Water 1, Water 2, and hexadecane.

**Video S3. Coordinated handling of water and ethanol as a miscible LCL by the P2PV-HPR.** The 3D-printed CC executes the sequential delivery of three liquids: Water 1 (green), Water 2 (yellow), and Ethanol (orange), which serves as a miscible LCL. The loading procedure consists of four steps: (1) the water reservoirs are filled; (2) the main channel and air waste are filled with the HCL; (3) the control HCL is loaded into the P2PV-HPR inlet, forming a P2PV and trapping an air gap; and (4) ethanol is loaded. Upon connection of a paper pump, the liquids are sequentially released in the following order: Water 1, Water 2, and ethanol.

**Video S4. Multiphase MCR demonstration.** The 3D-printed CC encoding the multiphase MCR sequentially delivers four liquids including Water 1 (green), Water 2 (yellow), Ethanol (orange) and Hexadecane (red). The water reservoirs are first loaded, and the main channel and the air wastes are filled with the HCL. The control HCL is then filled to form MDVs and P2PVs while trapping air gaps, followed by loading of ethanol and hexadecane, which are miscible and immiscible LCLs, respectively. Upon connection of a paper pump, the MCR enables sequential release of the four liquids into the main channel through conditional propagation of liquid events, whereby complete drainage of a reservoir establishes an air connection to the next, triggering its release.

**Video S5. Multiphase MCR CC for LNP fabrication.** A self-contained LNP fabrication process is performed by an autonomous 3D-printed MCR CC without peripherals. Buffer, stock RNA solution, lipid solution in ethanol, and hexadecane are loaded into their respective reservoirs, and upon actuation by paper-pump connection, the CC autonomously dilutes the stock RNA in buffer and mixes it with lipids in ethanol for RNA-LNP fabrication using integrated micromixers, followed by collection of the assembled LNPs into the integrated collection chamber for retrieval via displacement by immiscible hexadecane.
